# Force-field perturbations and muscle vibration strengthen stability-related foot placement responses during steady-state gait

**DOI:** 10.1101/2023.10.13.562176

**Authors:** A.M. van Leeuwen, S.M. Bruijn, J.C. Dean

## Abstract

Mediolateral gait stability can be maintained by coordinating our foot placement with respect to the center-of-mass (CoM) kinematic state. Neurological impairments can reduce the degree of foot placement control. For individuals with such impairments, interventions that could improve foot placement control could thus contribute to improved gait stability. In this study we aimed to better understand two potential interventions, by investigating their effect in neurologically intact individuals. The degree of foot placement control can be quantified based on a foot placement model, in which the CoM position and velocity during swing predict subsequent foot placement. Previously, perturbing foot placement with a force-field resulted in an enhanced degree of foot placement control as an after-effect. Moreover, muscle vibration enhanced the degree of foot placement control through sensory augmentation whilst the vibration was applied. Here, we replicated these two findings and further investigated whether Q1) sensory augmentation leads to an after-effect and Q2) whether combining sensory augmentation with force-field perturbations leads to a larger after-effect, as compared to force-field perturbations only. In addition, we evaluated several potential contributors to the degree of foot placement control, by considering foot placement errors, CoM variability and the CoM position gain (β_pos_) of the foot placement model, next to the R^2^ measure as the degree of foot placement control. Sensory augmentation led to a higher degree of foot placement control as an after-effect (Q1). However, combining sensory augmentation and force-field perturbations did not lead to a larger after-effect, as compared to following force-field perturbations only (Q2). Furthermore, we showed that, the improved degree of foot placement control following force-field perturbations and during/following muscle vibration, did not reflect diminished foot placement errors. Rather, participants demonstrated a stronger active response (higher β_pos_) as well as higher CoM variability.

## Introduction

Controlling where we place our feet mediolaterally from our center-of-mass (CoM) ensures stability during (steady-state) walking [1, 2]. Healthy individuals actively steer their swing leg [3, 4] to enact this mechanism [5, 6]. However, in older adults and individuals with neurological impairments the coordination of the CoM kinematic state (during swing) and subsequent foot placement can be diminished [7–9].

For stroke patients classified as at high fall risk, the coordination of CoM kinematic state and foot placement is clearly disrupted, whereas stroke patients at low fall risk still coordinate foot placement similarly to healthy controls [9]. Thus, training the relationship between foot placement and CoM kinematic state (hereafter referred to as “the degree of foot placement control”) may improve mediolateral gait stability and prevent falls. A linear model has been demonstrated to describe the relationship between the CoM kinematic state during swing and subsequent foot placement [5]. Since perturbing sensory information elicits responses predictably according to this model [7], we consider this linear foot placement model to reflect an internal model, defining target foot placement. That the target foot placement defined by this control strategy is aimed at maintaining gait stability followed from external lateral stabilization studies, showing less foot placement control when externally stabilized [10, 11]. Measures of the explained variance of this model, such as R^2^ and partial correlation measures can as such be considered “the degree of foot placement control” with respect to the CoM. In turn, the unexplained variance of the foot placement model (i.e. the residuals) can be considered as “foot placement errors”; emerging from motor noise, loose control or sensorimotor impairments.

The degree of foot placement control has been shown to increase as an after-effect following training sessions within a perturbing force-field, both in stroke patients [12], and in neurologically intact individuals [13, 14]. Even in neurologically intact individuals, CoM variations are not entirely accounted for by subsequent modulation in foot placement [5]. Foot placement errors persist, which are only partially accounted for by ankle moment control [15]. It thus seems that the remainder of these foot placement errors, is small enough for healthy, young individuals to remain stable. However, a perturbing force-field may increase the foot placement errors to such an extent that even healthy individuals are driven to tighten their foot placement control to minimize the error. Consequently, error-based learning [16] may underlie the adaptations to a perturbing force-field [13, 14]. The force-field perturbs foot placement, as such changing the outcome of the motor plan actuating muscles to steer the swing leg towards the target location. Assuming we perceive the resulting foot placement errors as a motor execution error, rather than having aimed for a wrong target foot placement, participants would adapt this motor plan to improve foot placement control [16].

Furthermore, the degree of foot placement control can be improved whilst walking with augmented proprioception, provided through muscle vibration [17]. Applying muscle vibration to either the stance leg or the swing leg hip adductors, caused more medial or more lateral foot placement, respectively [17]. Controlling vibration based on step-by-step pelvis displacement (as a proxy for step-by-step CoM displacement) strengthened the relationship between the CoM kinematic state and foot placement [5]. These improvements in foot placement control have been attributed to sensory augmentation. In other words, the ratio between detected relevant sensory information, in this case the CoM kinematic state (stance leg vibration) or the swing leg’s mechanical state (swing leg vibration), and sensory noise is thought to be improved by the timed muscle vibration. Consequently, as a result of better sensory detection, the degree of foot placement control increased whilst the muscle vibration was active [17].

It remains to be investigated whether sensory augmentation has long lasting effects. Since timed muscle vibration is thought to improve sensory detection of the CoM kinematic state, as well as swing leg mechanical state, we considered it unlikely that sensory augmentation by itself would induce any after-effects. Improved foot placement control due to sensory augmentation would require the vibration to be active. To test the assumption that sensory augmentation only affects foot placement control whilst muscle vibration is applied, we tested Q1) whether sensory augmentation leads to an after-effect.

If sensory augmentation does lead to an after-effect, it would prove our assumption outlined above wrong. It would mean that apart from an immediate sensory augmentation effect, muscle vibration may facilitate motor adaptation. Perhaps, an improved sensory signal to noise ratio, may allow for a more accurate update of the motor plan, following better detection of erroneous motor execution of foot placement.

If so, sensory augmentation may especially enhance error-based motor learning, when applied during foot placement perturbations. To investigate this possibility, we tested Q2) whether adding sensory augmentation to perturbing force-field training enhances its positive effects on the degree of foot placement control.

In addition, we replicated that R1) a perturbing force field results in an increased degree of foot placement control as an after-effect and R2) that augmented proprioception enhances the degree of foot placement control during muscle vibration. Lastly, we checked whether, in line with our assumptions regarding error-based learning, force field perturbations increased foot placement errors.

## Methods

The research proposal and hypotheses have been preregistered on OSF: https://doi.org/10.17605/OSF.IO/FDCHU. We have deviated from the preregistered research proposal, by changing our introduction, to improve focus and clarity of the manuscript. In addition, we have checked one of our assumptions, namely that the force-field would induce larger foot placement errors, through an additional statistical test. Furthermore, although we retained our preregistered main outcome measure, we now support this main outcome measure with different secondary outcome measures than preregistered. In doing so, we could gain better insight into what changes in our main outcome measure reflected.

### Participants

Neurologically intact adults (18-40 years old) were recruited. The final sample size was partly determined by a Bayesian sequential sampling approach (i.e. sampling until a threshold of meaningful evidence was reached) and partly by collecting sufficient data to explore order effects of the three training types (see supplementary material A). We initially aimed for recruiting 18 participants (i.e. 3 participants for each order of training types), yet planned to continue to add participants until we reached a BF_10_ of 10 or a BF_01_ of 0.1 (indicative of strong relative evidence respectively for the alternative or the null hypothesis) or a preset maximum of 24 participants. As preregistered, the sequential sampling approach was based on the Bayes Factors of both R1 and R2.

Since, for our main outcome measure, we did not reach a Bayes Factor indicating strong relative evidence for R1 and R2, we ended up with the predetermined maximum of 24 participants (see table 1 for participant characteristics). All participants signed an informed consent form prior to participating in the experiment, and ethical approval was granted by the Institutional Review Board of the Medical University of South Carolina.

**Table 1.** Participant characteristics.

| **Participant characteristics** |  |
| --- | --- |
| **Gender** | 17 female 7 male |
| **Age** | $26 \pm 3.7$ SD years old |
| **Height** | $170 \pm 9.8$ cm |
| **Weight** | $74 \pm 15$ kg |

### Experimental protocol

Participants visited the lab for one session, in which they were asked to walk on a treadmill at a speed of 1.2 m/s, in one familiarization and three training conditions. During all walking conditions, participants wore a harness and were instructed to look forward as much as possible. As a safety precaution, one experimenter was always standing next to the treadmill, chatting to them and making sure they were comfortable throughout the experiment.

During familiarization participants walked for ten minutes in a transparent force-field set-up, (i.e. the leg cuffs of the force-field were mounted, but the force-field produced minimal force on the legs). During familiarization no sensory augmentation was applied. In the training conditions participants walked an eight-minute episode either in a perturbing force-field, with sensory augmentation or with sensory augmentation whilst in the perturbing force field. Before and after each training episode participants walked for two minutes again within a transparent force field and without sensory augmentation.

**Figure 1.**
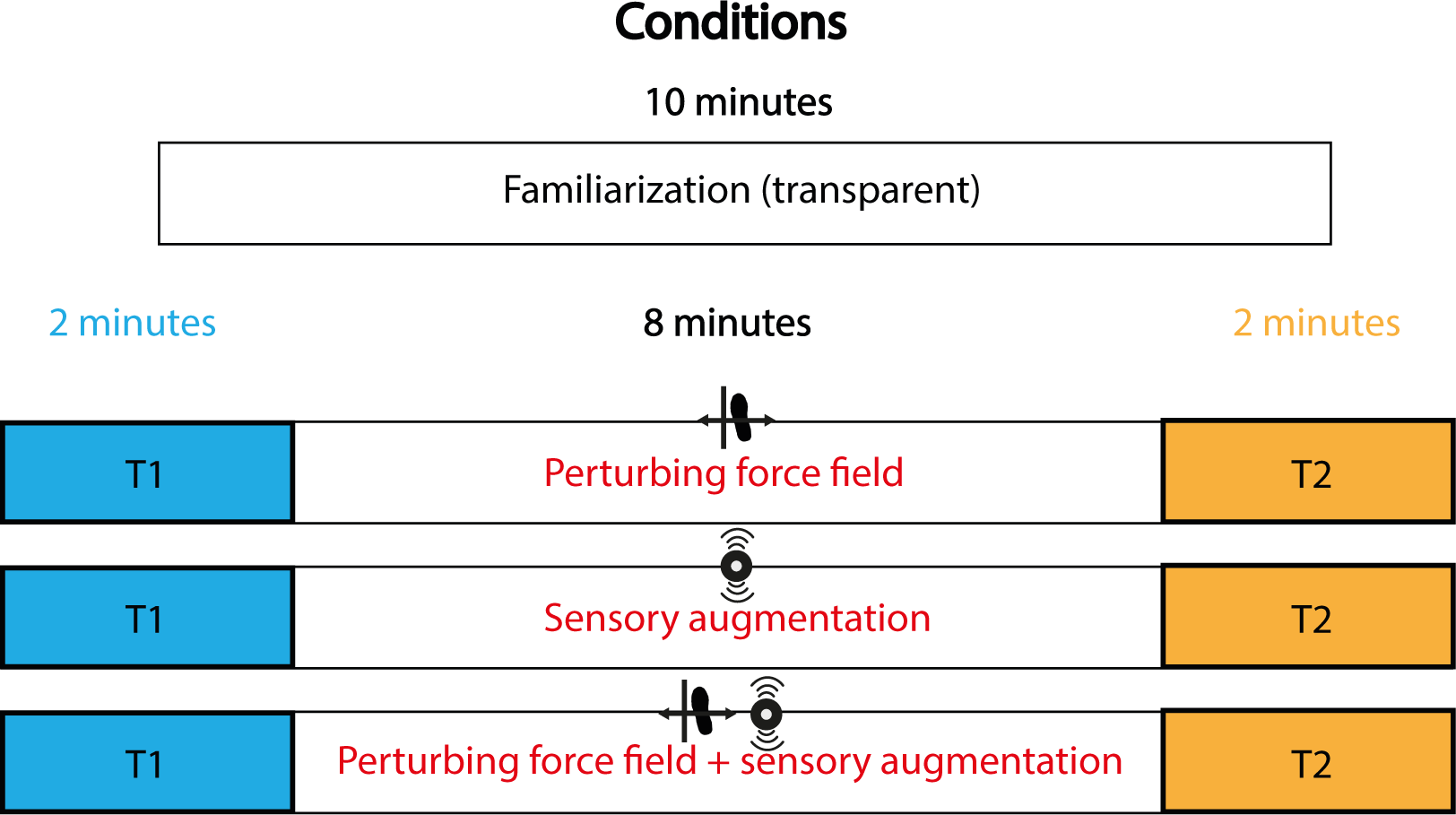
Walking conditions for the participants. After familiarization participants walked in three training conditions, in a randomized order. In Condition 1 the training encompasses walking in a perturbing force-field. In Condition 2, participants walked whilst receiving sensory augmentation. In Condition 3 the perturbing force-field training was combined with sensory augmentation. Training episodes lasted 8 minutes. Each training episode was preceded by and concluded with a transparent walking condition of 2 minutes, respectively T1 and T2, for which neither force-perturbation nor sensory augmentation was applied.

### Data collection and experimental set-up

#### Kinematics

Kinematics of the pelvis and feet (heel marker) were recorded using PhaseSpace, sampled at 120 Hz. Additional markers were placed on the leg-cuffs of the force field to operate the perturbations.

#### Force-field

The force-field used in the current study is described in more detail in [18]. Participants wore a cuff on both calves, with two bars on the lateral side of each cuff for the force-field wire to run through. The force-field wires were attached to sliders in the front and back of the treadmill, which, during the trial were actuated to slide in mediolateral direction, and as such perturb foot placement in the mediolateral direction. Prior to the walking conditions, we aimed to tension this wire until the same value for each participant (180 N/m), which was earlier determined to be an appropriate stiffness to induce clear effects on step width [13]. The force-field’s responses were triggered by the heel markers (phase of the gait cycle) and the perturbation magnitude was determined based on how far the pelvis marker deviated from its median mediolateral displacement, as computed for the final 50 strides of T1.

In this way, the perturbing force-field was controlled based on individual stepping characteristics. The perturbations aimed at pushing the leg towards taking a wide step when a narrow step was warranted to maintain gait stability, and vice-versa. For the final 50 steps of the T1 period, we quantified the step width and the mediolateral displacement of the pelvis from the stance foot at the start of the step (defined as the instance where the heel marker switched from moving posterior to moving anterior [19]). We then sorted the step width values in descending order and the pelvis displacement values in ascending order, and calculated the best-fit linear relationship between the sorted variables. This relationship is the basis for our force-field control. It perturbs the normal positive correlation between pelvis displacement and step width, whilst ensuring that participant step widths approximately spanned their normal range. For example, if for a given step, the pelvis displacement is at the very low end of the range seen in this participant, the force-field will push the swing leg toward a mediolateral location corresponding to a step width at the high end of the range in this participant and vice versa. If the pelvis displacement is around the median value for a participant however, the force-field will push the swing leg toward approximately the median step width value for this participant. Thus, the farther the pelvis is from the median pelvis displacement in T1, the larger the perturbation magnitude will be.

#### Sensory augmentation

Sensory augmentation was achieved by applying muscle vibration using tactors as in [17]. A tactor was placed across each m. gluteus medius, i.e. between the trochanter major and the iliac crest. The tactors were triggered based on the pelvis displacement at the start of the step. Depending on whether this pelvis displacement was smaller or larger than the median pelvis displacement during the final 50 steps of the corresponding T1 episode, either the stance or the swing leg muscle vibrator was activated during that step. The further from the median pelvis displacement, the higher the vibration frequency (ranging between 34 and 74 Hz). The vibration was turned on at the start of the step and lasted for a duration of the mean step period during the final 50 steps of T1.

### Data processing

#### Gait event detection

For the data analysis, gait event detection was based on the heel markers. Right and left heel-strikes were defined as minima in vertical position of the right and left heel marker. Right and left toe-off events were defined as the maxima in vertical velocity of respectively the right and left heel markers.

#### Outcome measures

Our main outcome measure is the relative explained variance (R^2^) of the foot placement model (equation [1]). This model was fit for each individual, over 50 stride (i.e. over 50 left and 50 right steps) blocks. The R^2^ was considered as “the degree of foot placement control”, indicative of how well foot placement is predicted by the CoM kinematic state (ML*_CoM_* & ML*_CoMvel_* at terminal swing). The measured pelvis kinematic state served as a proxy for the CoM kinematic state. *FP* was defined as the demeaned distance between the pelvis and the mediolateral placement of the swing foot at the end of the step. β_pos_ and β_vel_ are the regression coefficients and ε is the residual of the model.

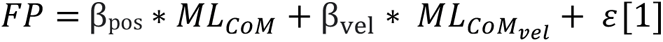

As our main outcome measure, we based our conclusions on the statistical tests on R^2^ for CoM kinematic state at the end of the step. This measure reflects the degree to which active and passive foot placement mechanisms achieved stability-related foot placement goals.

To provide more insight into what underlies the potential changes in R^2^, we included several secondary outcome measures.

Changes in the R^2^ can reflect changes in the standard deviation of the residual (foot placement error), β_pos_, β_vel_ or in the CoM kinematic state variance (see figure 2). In order to interpret the changes in R^2^ we included foot placement error, β_pos_ and the variability in mediolateral CoM position as secondary outcome measures to the results.

**Figure 2.**
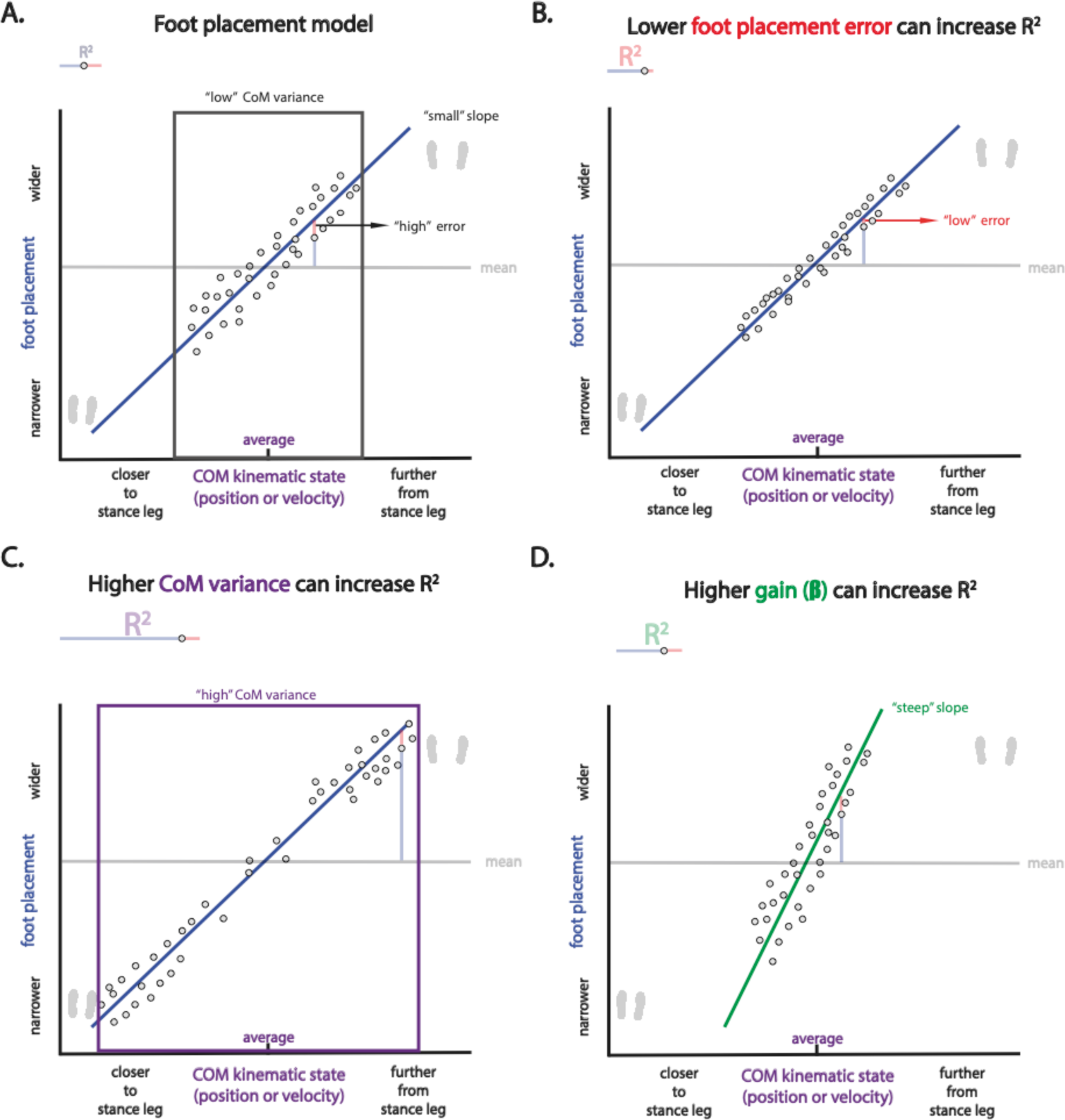
The degree of foot placement control can increase in different ways. Fictive data to illustrate that a change in R^2^ can reflect different types of adaptation. Figure from [20].

In previous work, SW (i.e. the demeaned mediolateral distance between the left and right foot, both during stance) has often been considered as the dependent variable of the foot placement model (equation [2]) [4, 5, 10, 15, 21]. In the current study we opted for FP as the dependent variable (equation [1]), since this measure reflects more directly the stability driven goal of placing the foot at an appropriate distance from the CoM. The results based on SW can be found in the supplementary material (Supplementary material B).

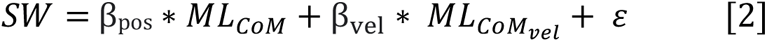

#### Statistics

We used four Bayesian equivalents of paired t-tests, to replicate two earlier findings (R1&R2) as well as to test two novel research questions (Q1 & Q2). For R1&R2 we used a one-sided t-tests as we had clear expectations of the direction of the change. For Q1 & Q2 we used a two-sided t-test.

The Bayesian t-tests were performed using the default’s prior in JASP (Cauchy centered around 0 with a 0.707 scale).

To test R1, that a perturbing force field induces an increased degree of foot placement control as an after-effect, we tested the last 50 steps of T1 of Condition 1 against the first 50 steps of T2 of Condition 1. We expected an increased degree of foot placement control at the start of T2 as compared to the end of T1.

To test R2, that sensory augmentation enhances the degree of foot placement control during muscle vibration, we tested the last 50 steps of T1 of Condition 2 against the whole training episode of Condition 2 (averaging over the outcome measures separately computed for each 50-stride block). We expected an increased degree of foot placement control during the training episode as compared to T1.

To test Q1, whether sensory augmentation leads to an after-effect on the degree of foot placement control, we tested whether augmented proprioception caused any difference between the first and last transparent episode of Condition 2 (i.e. the last 50 steps of T1 and the first 50 steps of T2 of condition 3).

To test Q2, whether adding muscle vibration to perturbing force-field training enhances its positive effect on the degree of foot placement control, we tested whether the change in the degree of foot placement control between T2 and T1 is different in Condition 1 as compared to Condition 3. Thus, prior to the statistical analysis, we computed ΔR^2^ for each condition, by subtracting the R^2^ from the last 50 steps of T1 from the R^2^ from the first 50 steps of T2.

In the results we report the Bayes Factors (BF) as well as the degree of evidence (i.e. anecdotal/moderate/strong/very strong/extreme) based on [22].

The data and analyses can be found on surfdrive: https://surfdrive.surf.nl/files/index.php/s/T10VlkDR9SH8fxs, and will be published on Zenodo if the manuscript is accepted for publication.

## Results

Below we discuss the results based on Model 1, i.e. the model with foot placement (FP) as dependent variable. In supplementary material B, we present the results for Model 2, i.e. the model with step width (SW) as the dependent variable.

### Perturbation check

Force field perturbations were effective in increasing foot placement error (Figure 3), indicating the force field induced perturbations in line with our expectations. This was supported by extreme evidence (BF_10_ = 36800).

**Figure 3.**
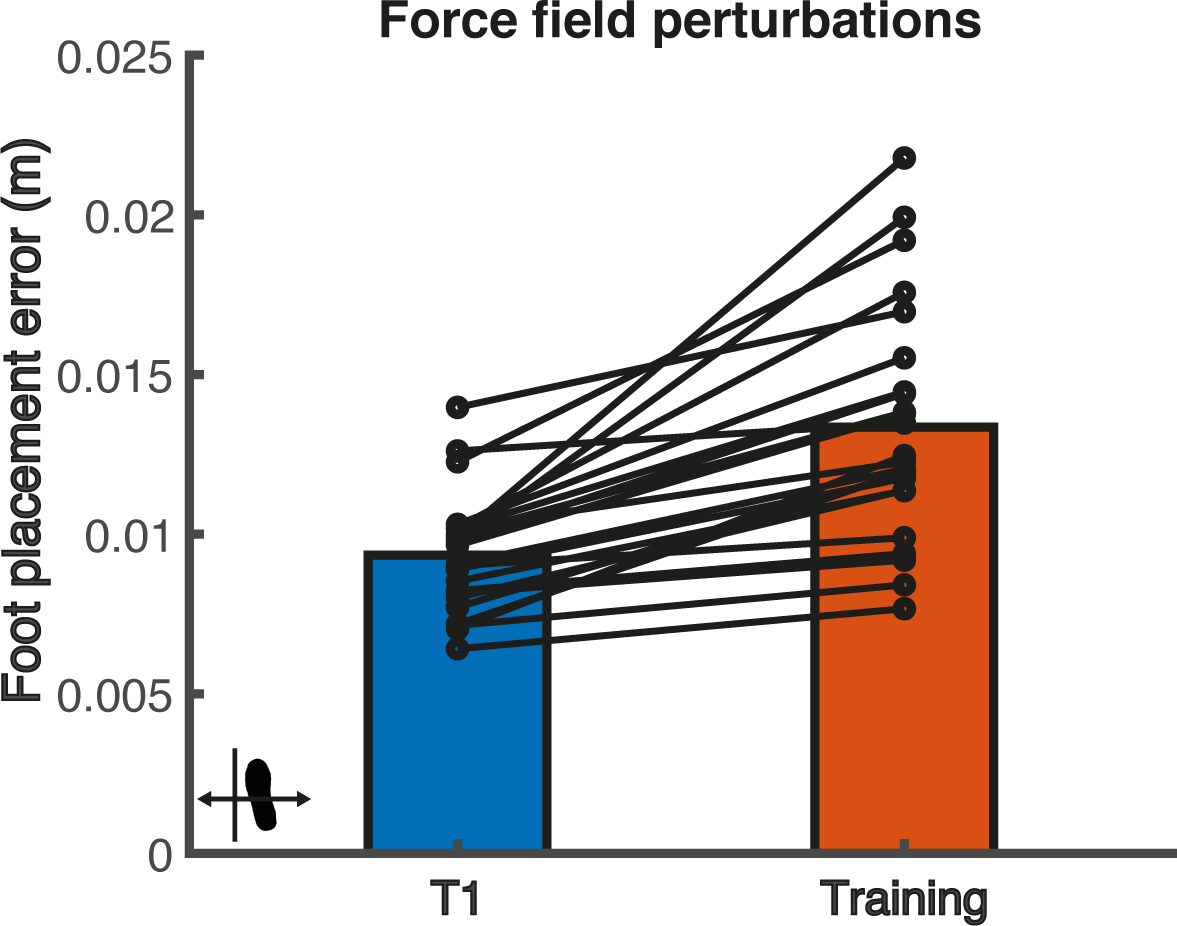
Foot placement errors before (T1) and during (Training) force field perturbations. The black circles indicate individual data points. The magnitude of foot placement errors was determined as the standard deviation of the residuals of the foot placement model with FP as the dependent variable and CoM kinematic state predictors at terminal swing.

### Replication

#### R1

We found strong evidence (BF_+0_ = 19.42) that a perturbing force field results in an increased degree of foot placement control as an after-effect (R1, Figure 4).

**Figure 4.**
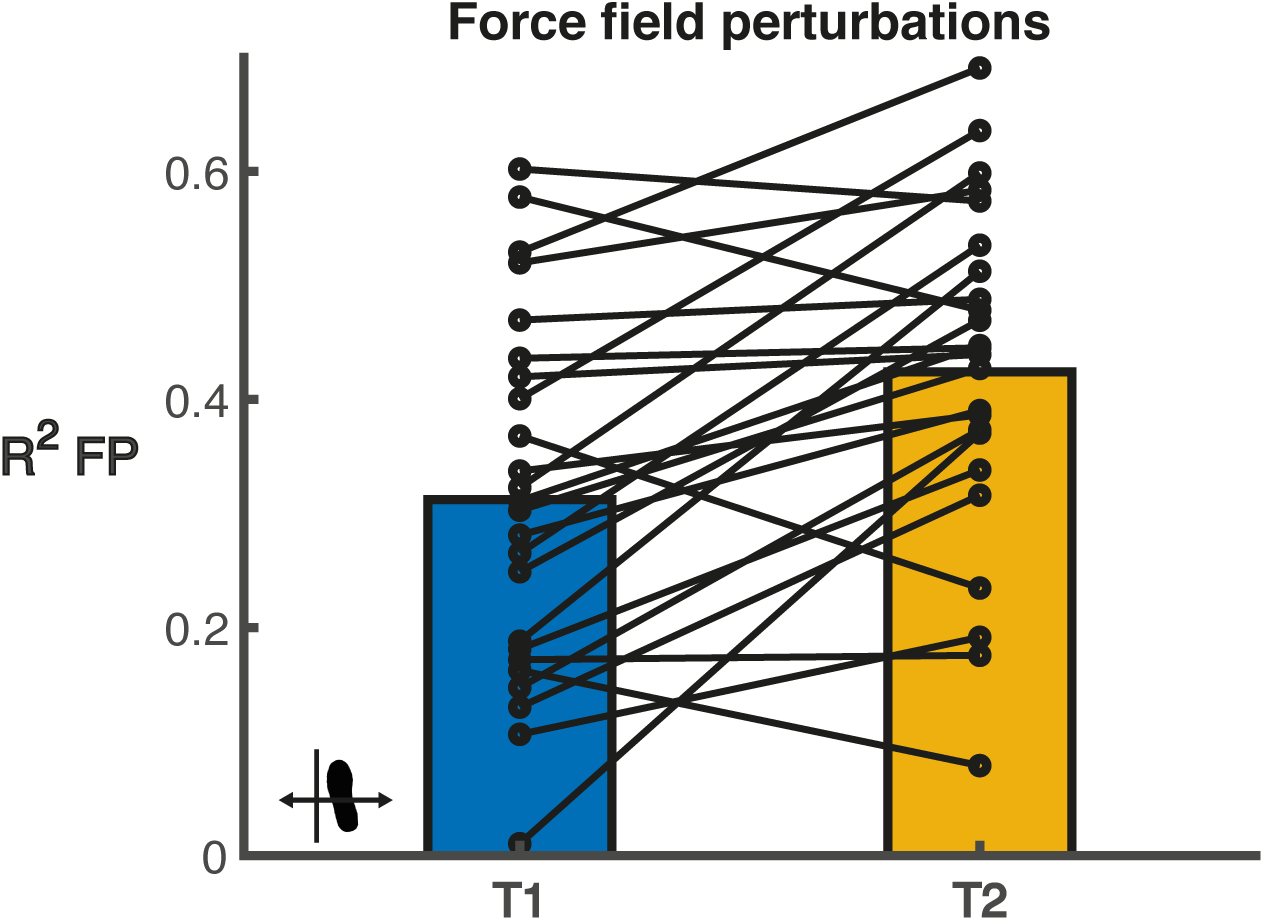
The degree of foot placement control (R^2^) before (T1) and after (T2) force field perturbations. The black circles indicate individual data points. R^2^ was determined from the foot placement model with FP as the dependent variable and CoM kinematic state predictors at terminal swing.

Foot placement error (extreme evidence, BF_10_ = 143), variability in mediolateral CoM position (moderate evidence, BF_10_ = 9.83) and the regression coefficient of ML*_CoM_* (very strong evidence, BF_10_ = 90.58) all increased as an after-effect.

**Figure 5.**
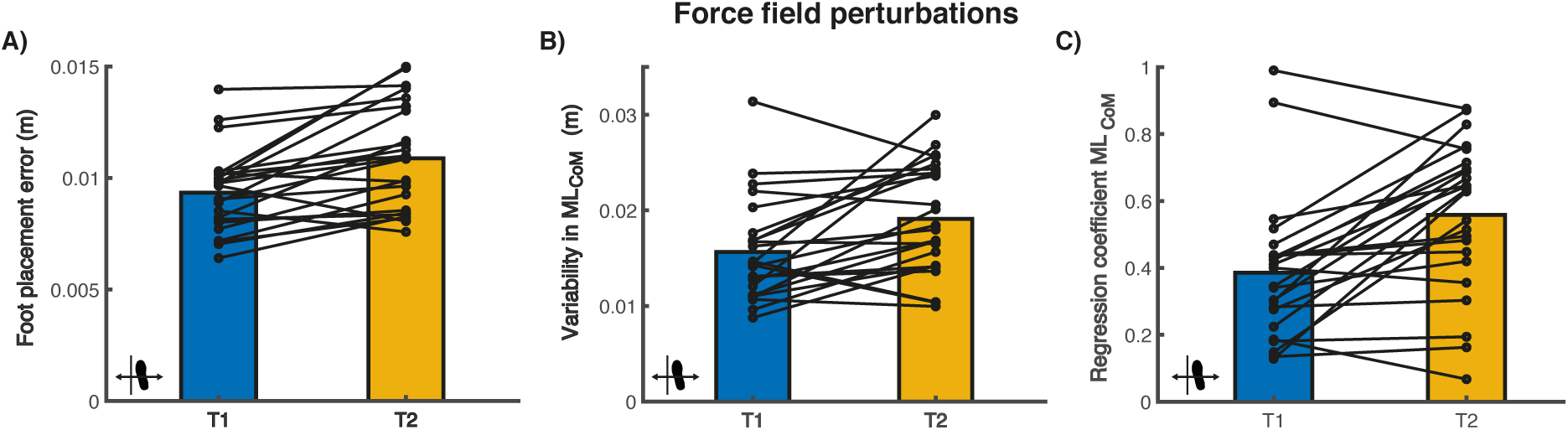
Foot placement error (Panel A), mediolateral CoM position variability (panel B) and β_pos_ (panel C) before (T1) and after (T2) force field perturbations. The black circles indicate individual data points. These secondary outcomes provide more insight into the underlying foot placement control changes reflected in the R^2^ of figure 4.

#### R2

We found extreme evidence, (BF_+0_ = 57700) that augmented proprioception enhances the degree of foot placement control during the muscle vibration (R2, Figure 6).

**Figure 6.**
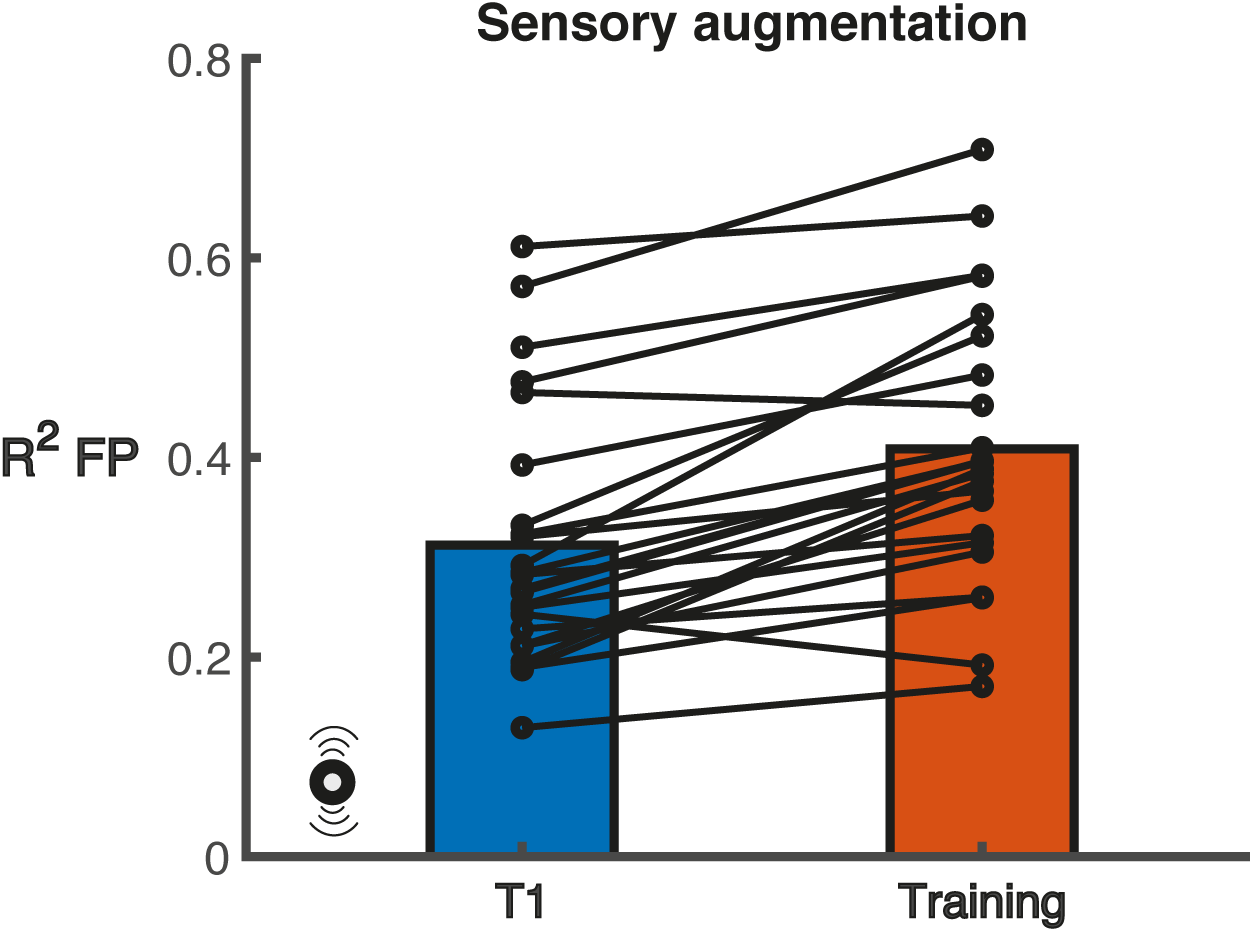
The degree of foot placement control (R^2^) before (T1) and during (Training) sensory augmentation. The black circles indicate individual data points. R^2^ was determined from the foot placement model with FP as the dependent variable and CoM kinematic state predictors at terminal swing.

Anecdotal evidence was found supporting an increase in foot placement error during sensory augmentation (BF_10_ = 1.14). The variability in mediolateral CoM position (very strong evidence, BF_10_ = 36.06) as well as the regression coefficient β_pos_ (extreme evidence, BF_10_ = 1613.86) increased during sensory augmentation.

**Figure 7.**
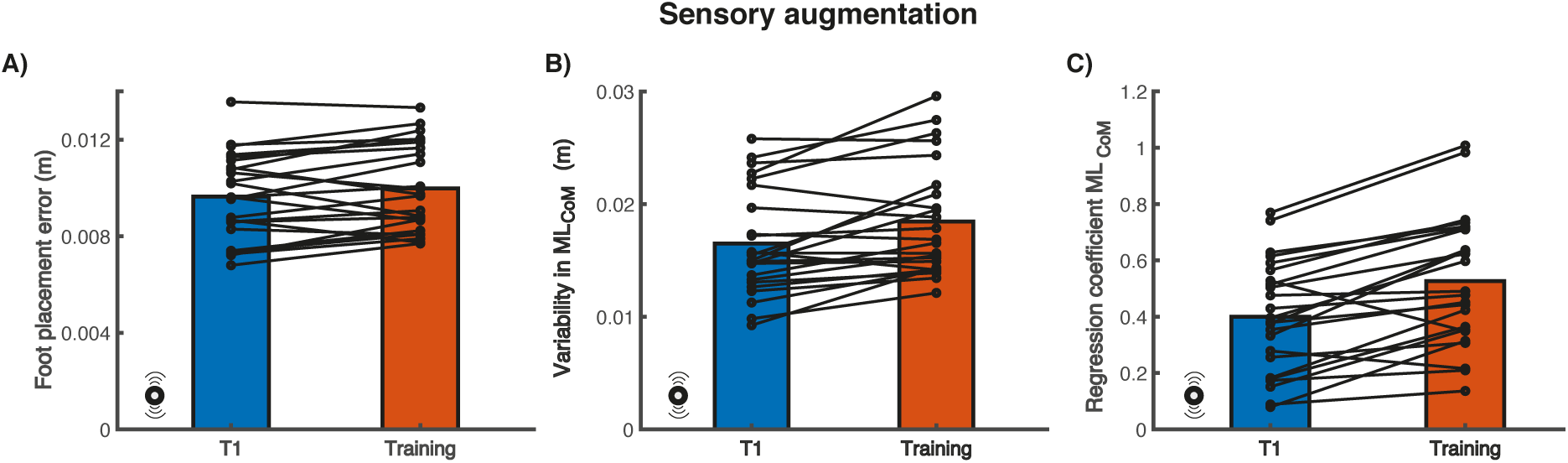
Foot placement error (Panel A), mediolateral CoM position variability (panel B) and β_pos_ (panel C) before (T1) and during (Training) sensory augmentation. The black circles indicate individual data points. These secondary outcomes provide more insight into the underlying foot placement control changes reflected in the R^2^ of figure 6.

### Sensory augmentation after-effect?

We found strong evidence (BF_10_ =13.96) that sensory augmentation caused an after-effect (Q1). In Figure 8 it can be seen that the degree of foot placement control increased in T2 as compared to T1.

**Figure 8.**
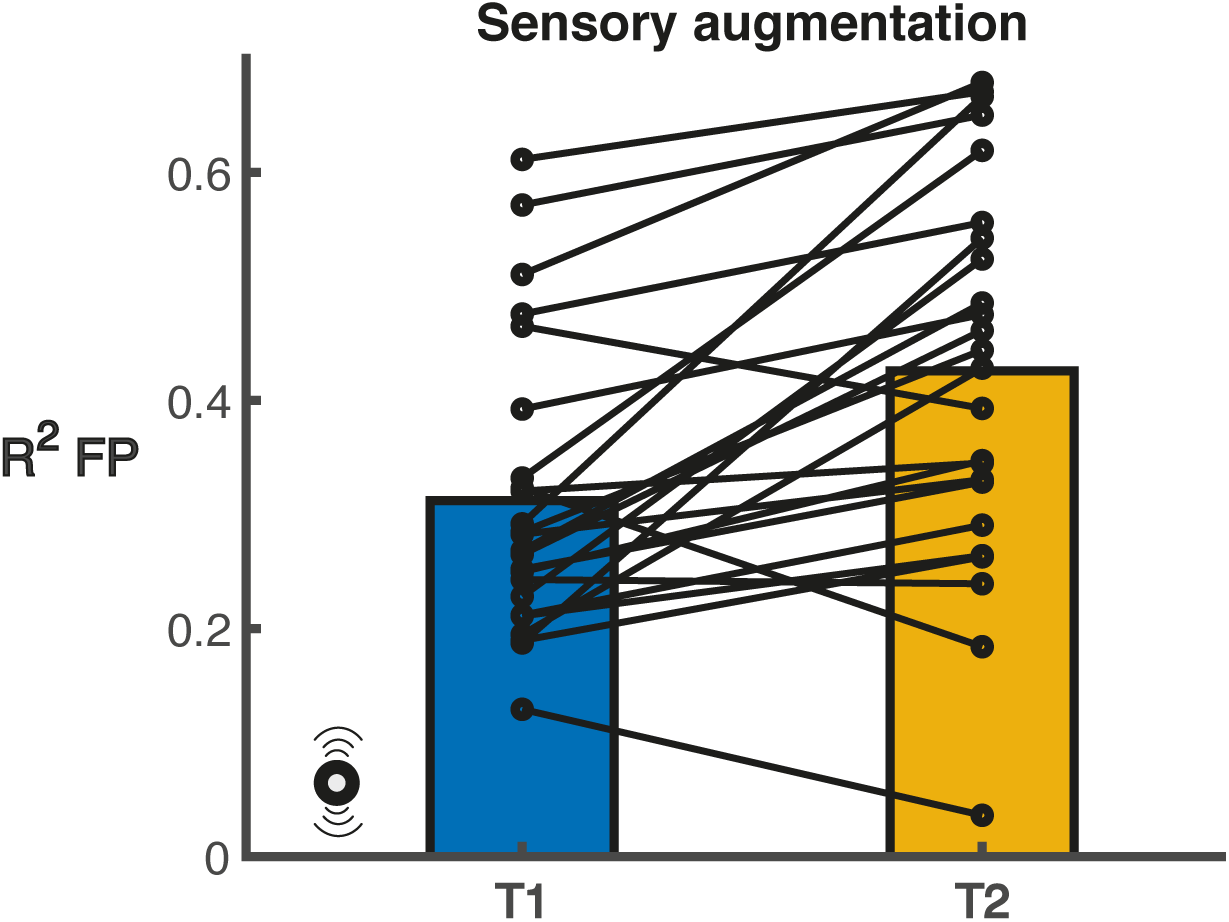
The degree of foot placement control (R^2^) before (T1) and after (T2) sensory augmentation. The black circles indicate individual data points. R^2^ was determined from the foot placement model with FP as the dependent variable and CoM kinematic state predictors at terminal swing.

We found anecdotal evidence against an after-effect in foot placement error following sensory augmentation (BF_10_ = 0.66). Variability in mediolateral CoM position increased as an after-effect of sensory augmentation (moderate evidence, BF_10_ = 5.80). Moreover, there was anecdotal evidence supporting an increase in regression coefficient β_pos_ as an after-effect (BF_10_ = 2.94).

**Figure 9.**
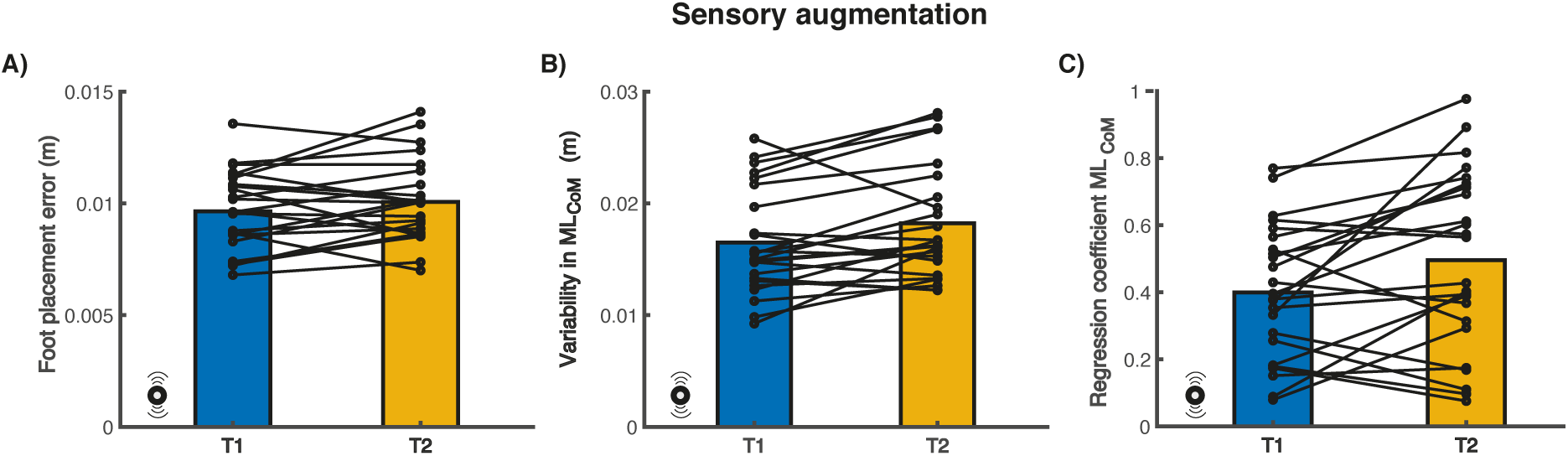
Foot placement error (Panel A), mediolateral CoM position variability (panel B) and β_pos_ (panel C) before (T1) and after (T2) sensory augmentation. The black circles indicate individual data points. These secondary outcomes provide more insight into the underlying foot placement control changes reflected in the R^2^ of figure 8.

### Force-field perturbations and sensory augmentation

We found moderate evidence (BF_10_ = 0.22) against a difference in after-effect magnitude after combining force-field perturbations and sensory augmentation as compared to after force-field perturbations only.

**Figure 10.**
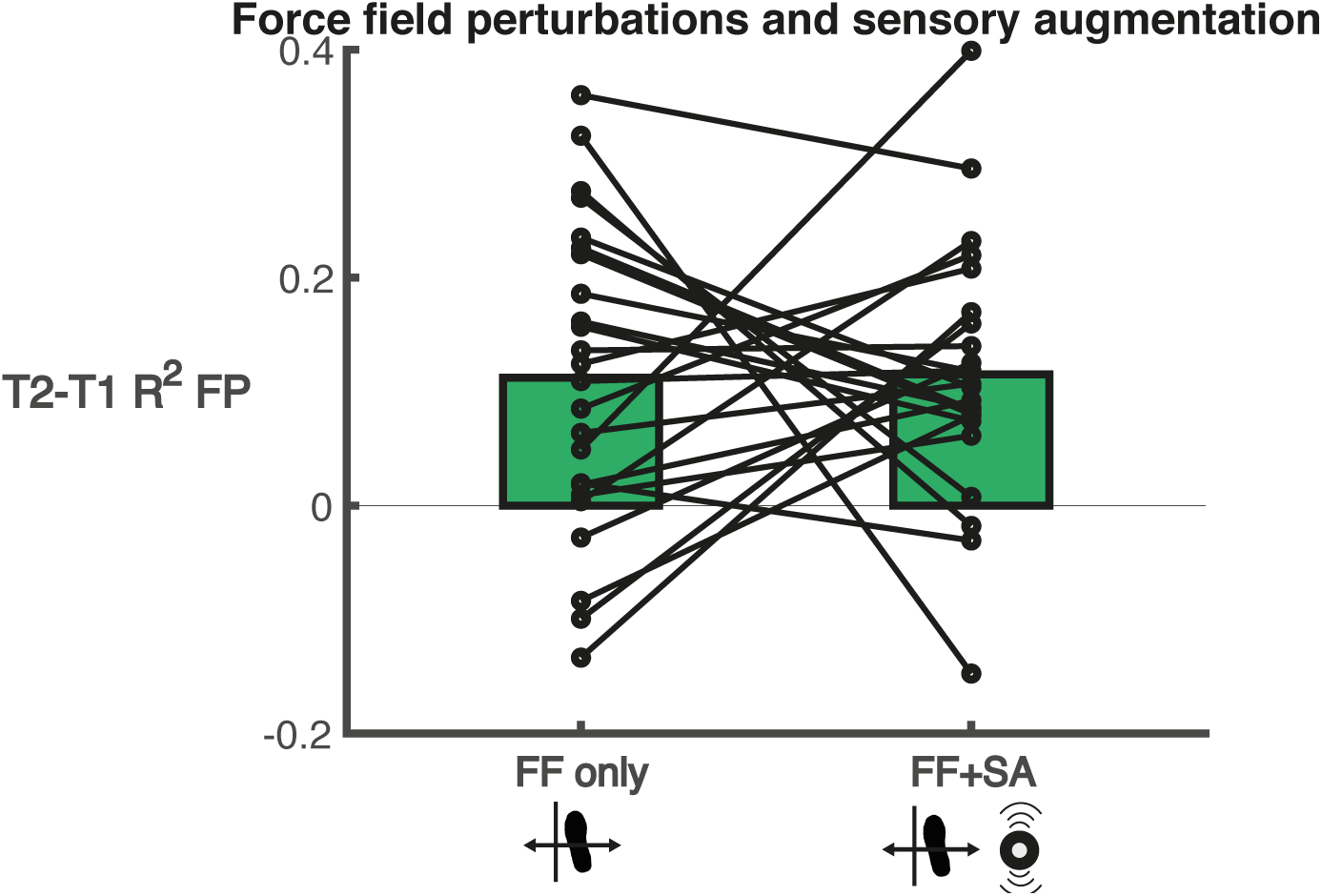
The change in the degree of foot placement control (R^2^) between T2 and T1 after force-field perturbations only and after a combination of force-field perturbations and sensory augmentation. The black circles indicate individual data points. R^2^ was determined from the foot placement model with FP as the dependent variable and CoM kinematic state predictors at terminal swing.

Similarly, we found moderate evidence against a difference in after-effect magnitude after combining force-field perturbations and sensory augmentation as compared to after force-field perturbations only, for the foot placement error (BF_10_ = 0.22), the variability in mediolateral CoM position (BF_10_ = 0.25) and the regression coefficient β_pos_ (BF_10_ = 0.26).

**Figure 11.**
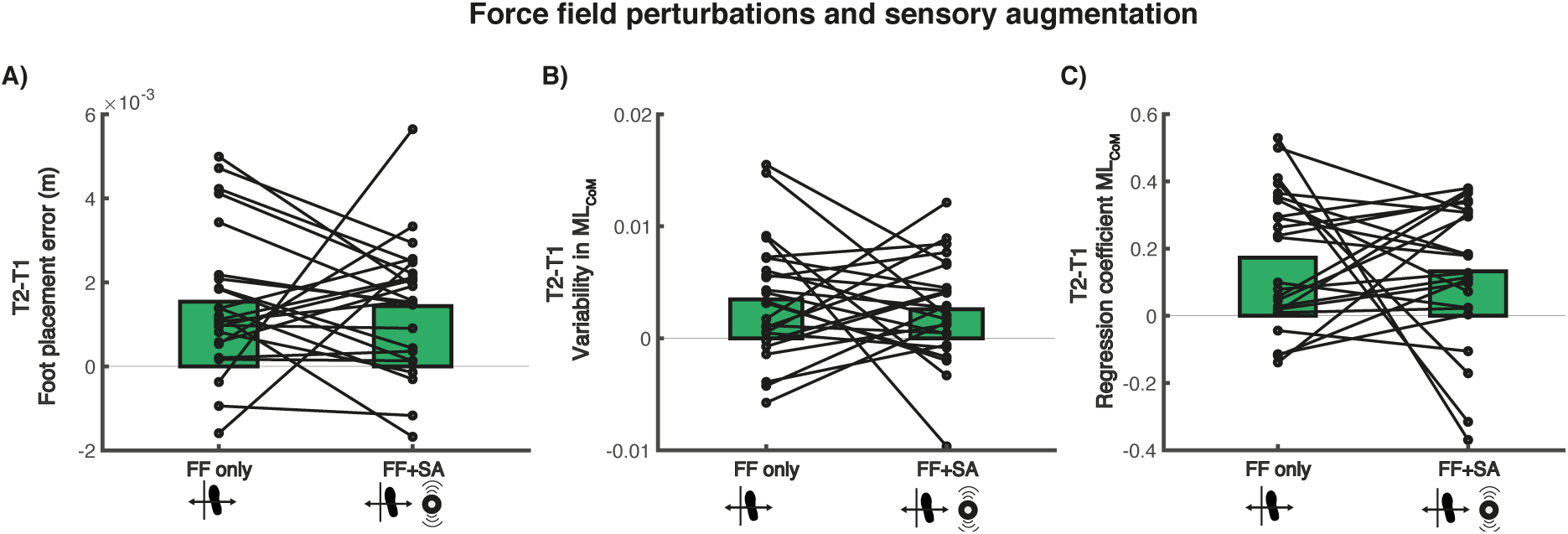
The change in foot placement error (Panel A), mediolateral CoM position variability (panel B) and β_pos_ (panel C), between T2 and T1 after force-field perturbations only and after a combination of force-field perturbations and sensory augmentation. The black circles indicate individual data points. These secondary outcomes provide more insight into the underlying foot placement control changes reflected in the delta R^2^ of figure 10.

## Discussion

By varying where we place our feet mediolaterally in accordance with variations in CoM kinematic state, we actively execute a stability control mechanism during steady-state walking [3, 4, 10]. Here, we replicated previous findings suggesting that such foot placement control can be enhanced, either through perturbation-based training [13, 14] or sensory augmentation using muscle vibration. Previous research suggests that improving mediolateral foot placement may help stroke patients [8, 9, 12] and older adults [7] to regain control over their gait stability. We aimed to better understand how (apparent) enhancements in foot placement control are achieved and tested whether combining two methods yielded larger training effects. Combining sensory augmentation and perturbation-based training did not result in augmented after-effects following perturbation-based training. Challenging our current understanding of its underlying mechanisms, sensory augmentation itself led to a higher degree of foot placement control as an after-effect.

### Perturbation induced error-based learning?

As expected, the force-field perturbations caused foot placement errors to increase whilst the perturbations were applied (Figure 3). Adapting to these errors could result in a gradual increase of the degree of foot placement control during perturbed walking, as was reported earlier [13], and as an after-effect, as was found before [13, 14] and replicated here (Figure 4). Although error-based learning [16] may have occurred, the current results show that the after-effect comes with an increase in foot placement errors (Figure 5 panel A). Thus, the adaptations to the force-field did not result in more precise foot placement once the perturbations were stopped. Instead, participants walked with a higher variation in their CoM kinematic state (figure 5, panel B) and with a higher gain (Figure 5, panel C) of the foot placement model (Model 1, β_pos_). It seems that participants adapted their motor plan to generate stronger active responses (as reflected by the higher gain), in response to deviations in their CoM kinematic state. This would favor the aim of placing the foot closer to the target position whilst the perturbations are still active. However, this adaptation will likely cause foot placement to overshoot a stable position once the perturbations stop. This can lead to larger deviations in CoM kinematic state. With a higher variation in CoM kinematic state, participants will have to place their feet at the wider end of the range that they usually walk at (Figure 2, Panel C), so that relatively their degree of foot placement control (R^2^) will improve, even though in absolute terms they stepped less precisely (higher foot placement errors).

### Replication: Sensory augmentation enhances the degree of foot placement control during muscle vibration

Improved sensory detection has demonstrated potential to enhance gait stability control [17, 23]. Muscle vibration studies showed that vibration of the hip abductor muscles causes responses in foot placement [7, 17], and when timed in accordance with deviations of the CoM kinematic state, can enhance the degree of foot placement control [17]. Here, we have replicated earlier findings showing an enhanced degree of foot placement control whilst the vibration is applied (Figure 6). Our secondary outcome measures provide more insight into what this improved degree of foot placement control reflects (Figure 7).

The degree of foot placement control did not improve as a result of a reduction in foot placement error magnitude (Figure 7, panel A). We found only anecdotal evidence to support a change in foot placement errors, and this change tended towards an increase as opposed to a decrease in foot placement error magnitude. Instead, a higher CoM variability (Figure 7, panel B), accompanied by a higher gain (Figure 7, Panel C), seem to underlie the higher degree of foot placement control. Based on the immediate effects only, this can be explained either in the context of the muscle vibration resulting in illusory sensory information, or in the context of muscle vibration improving the signal to noise ratio.

In the introduction, we proposed that timed muscle vibration, as sensory augmentation, may improve the sensory signal to noise ratio. However, perhaps the muscle vibration did not improve the signal to noise ratio, but rather created an illusion of a larger CoM deviation, or a more adducted swing leg respectively, that merely led to a stronger (higher gain) rather than to a more accurate response (no reduction in foot placement errors). In turn, these larger responses, based on “unreliable” sensory information, could have led to higher CoM variability.

Alternatively, the stronger response could be the result of sensory augmentation, improving the signal to noise ratio. With more certainty about the mechanical state, one may adopt less conservative foot placement control. With sensory augmentation, noise is less likely to result in a target position in the wrong direction and thus strong responses may pose less of a risk. More reliable sensory information due to sensory augmentation could therefore trigger recalibration of the internal model (Model 1) defining the targeted relationship between the CoM kinematic state during swing and subsequent foot placement. In this context, the increased CoM variability could characterize a gait pattern which is more dependent on foot placement control.

The latter explanation, i.e. in the context of sensory augmentation effects, would allow for better understanding of the after-effect we found following the training with muscle vibration.

### Sensory augmentation results in an after-effect

Against our initial assumption that muscle vibration had to be continuously applied for sensory augmentation to affect the gait pattern, after-effects were found following the timed muscle vibration. The degree of foot placement control increased (Figure 8), not due to a decrease in foot placement errors (Figure 9, panel A) but rather due to increased deviations in CoM mediolateral position (Figure 9, panel B) and its gain in Model 1(Figure 9, panel C).

Following the rationale outlined for the immediate sensory augmentation effects of muscle vibration, the changes in CoM variability and foot placement gain, may reflect a recalibrated internal foot placement model, as well as a gait pattern in which one chooses to rely more on foot placement control. The higher gain in the after-effect condition supports that the muscle vibration induced sensory augmentation rather than an unreliable illusion. If participants would have recalibrated the internal foot placement model (Model 1) due to a “perturbing” illusion, one would expect a lower gain in the after-effect condition. For example, an illusory more medial sensing of the CoM, will provoke too medial (i.e. with respect to the actual CoM position) foot placement, and participants would likely start to respond less strongly to the sensed (illusory) CoM kinematic state deviations. Thus, over time, resulting in a lower gain of Model 1. However, if participants were triggered to recalibrate their internal foot placement model due to sensory augmentation (i.e. more certainty of foot placement direction), the larger gain could be explained.

Since foot placement errors did not decrease, it is unlikely that the sensory augmentation improved error-based learning [16], as we had proposed earlier in the introduction. Rather, sensory augmentation may cause increased gains of the internal foot placement model (Model 1), if the sensory information can be considered more reliable. Since foot placement perturbations will lead to less predictable sensory outcomes, combining foot placement perturbations and sensory augmentation may mitigate sensory augmentation effects. Moreover, since sensory augmentation does not seem to improve error-based learning, it can be understood why we did not find an enhanced perturbation after-effect when adding muscle vibration to perturbation-based training.

### Combining force field perturbations and sensory augmentation does not lead to a larger after-effect

In Figures 10 and 11 it can be observed that combining the force field with sensory augmentation did not yield a larger positive after-effect as compared to force-field perturbations alone. As discussed above, although force-field perturbations seem to trigger error-based learning, sensory augmentation does not seem to affect the error-based learning process.

### Limitations

For the current study, a proper familiarization period was important, to avoid training effects to be confounded by familiarization effects. The ten minutes for familiarization were based on the recommendation of having at least six to seven minutes (or about 425 strides) for treadmill familiarization [24]. However, in the latter study, they did not test whether the specific outcome measures used in the current study, would stabilize after this time period. The degree of foot placement control has been found to change during a ten-minute familiarization period [21], and older adults still demonstrated changes in related outcome measures after multiple sessions of treadmill walking [20]. Yet, given the consistency of our findings with earlier work, as well as an absence of clear order effects (Supplementary material A), it is unlikely that the adaptation effects found reflect familiarization effects. Moreover, we ensured that each training condition had its own baseline (T1) trial, to limit the possibility of detecting effects due to walking duration only. Still, the absence of a control group, walking on the treadmill within a transparent force-field only, makes it that we cannot completely rule out effects of walking duration.

Another important factor we had to consider was to make sure that the training period was long enough to induce adaptation effects. Therefore, the eight-minute duration of the training episodes was based on earlier findings by [13], for which immediate effects of the force-field on the coordination between CoM kinematic state and foot placement appear to plateau after approximately eight minutes of force-field exposure. Since we replicated earlier findings of the effects of sensory augmentation and force-field perturbations, we may assume that, for the questions of this study, the training period was long enough.

By our choice of outcome measures we limited ourselves to a description of foot placement control. Although it has been established that foot placement control with respect to the CoM contributes to mediolateral gait stability [2, 10], we did not include a direct measure of gait stability, and can thus not make direct inferences on how force-field perturbations and sensory augmentation affect gait stability. Since the current study aims at giving better insight into what earlier found improvements in the degree of foot placement control [12, 17] reflect, as well as whether the effects of sensory augmentation and force-field perturbations could complement each other, further investigating the relationship with gait stability is beyond the scope of this study. Another study, has already established a link between foot placement outcome measures and clinical outcome measures [25]. Future work is needed to gain more insight into which (foot placement) outcome measures should improve for a gait stability training to be effective. Moreover, it remains to be investigated whether patient populations respond in the same way to these interventions, as the healthy adults in the current study. Future research should also focus on whether adaptation effects as measured on the treadmill translate to over ground walking.

Lastly, we admit that our current measures do not allow us to distinguish between adaptations to the internal foot placement model (Model 1), defining the target position, versus adaptations of the motor execution plan, defining the muscle activity to execute the foot placement response. For now, we speculate that the force-field operates at the level of the motor execution plan, likely leading to adaptations of the motor execution plan. Sensory augmentation on the other hand, may trigger adaptations of the internal foot placement model, updating desired target foot placement positions. Further experimental and modelling studies are required to untangle at which level the adaptations truly take place.

### Clinical perspectives

The training interventions studied here did not result in smaller foot placement errors, yet promoted stronger foot placement responses to deviations in CoM kinematic state. In stroke patients for whom foot placement control is disrupted to such an extent that the gains of the foot placement model (Model 1) tend to change sign [9], this may be a favorable adaptation. Perhaps, they can adapt to actively control their foot placement with a gain in the appropriate direction. However, for patients who aim to reduce CoM variability, such adaptations may be less desirable.

## Conclusion

We conclude that both sensory augmentation and force-field perturbations enhance the degree of foot placement control. Both methods result in a higher degree of foot placement control as an after-effect, but do not diminish foot placement errors. Instead, stronger active responses seem to be the result of both interventions. Combining the two interventions does not lead to augmented after-effects.

## Acknowledgements

The authors like to thank everyone that helped making the execution of this experiment possible in the short time that Moira visited Charleston. Especially, Heather Knight for preparing the measurement software, Keith Howard, for teaching how to operate the measurement systems, Camden Jacobs, for her encouragements and help in recruiting participants, Amelia Wright, for helping out during measurements, and of course everyone that participated in the experiment or asked someone else to do so.

## Supplementary material

### SA

#### Order effects?

**Figure SA 1.**
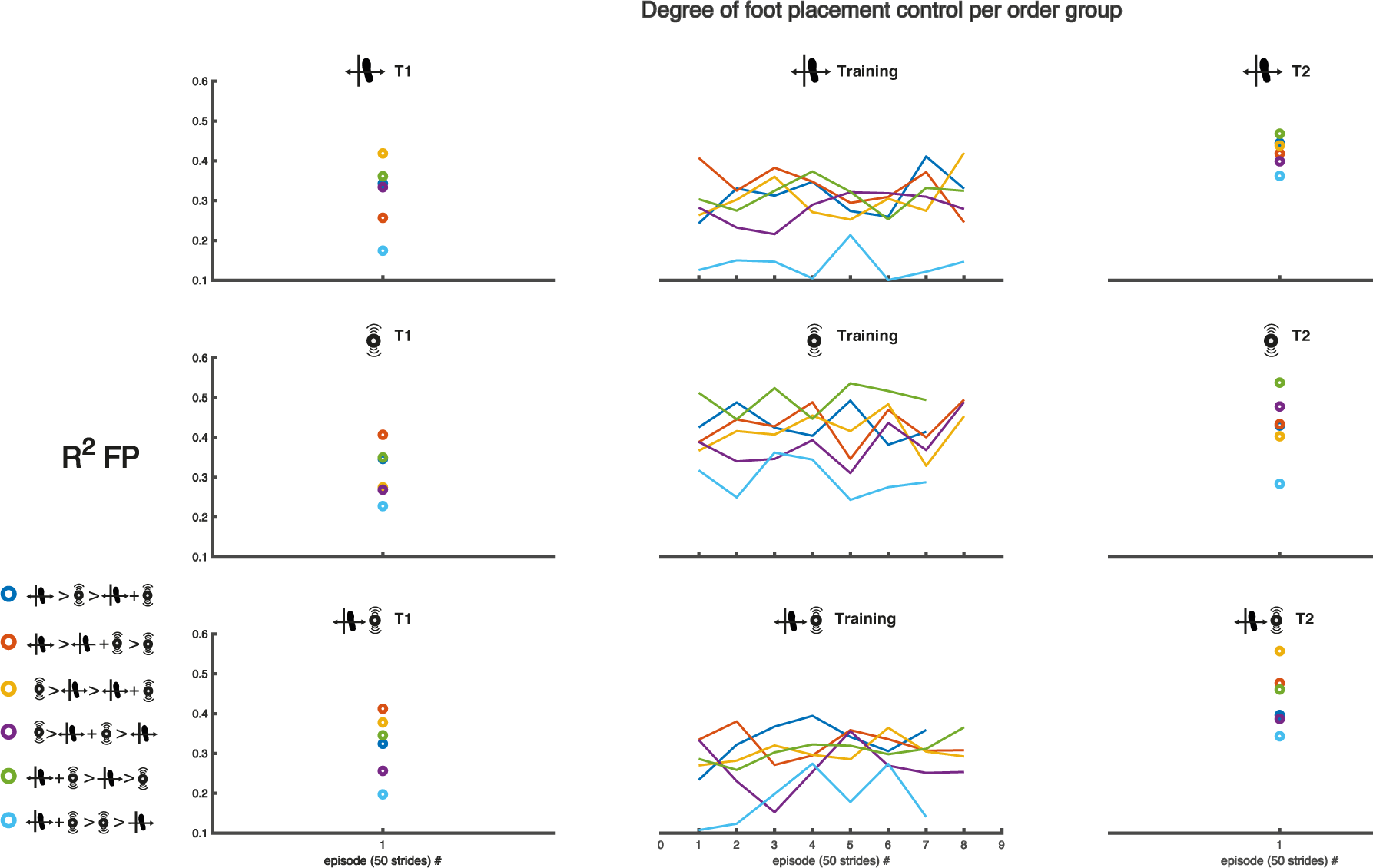
Exploring order effects. Participants were randomly assigned to six different possible orders of conditions. The figure shows the degree of foot placement control (for Model 1 with FP as the dependent variable) averaged over participants within each group. Based on visual inspection of this figure we concluded there were no clear order effects that may have affected our results.

### SB

#### Results with Model 2 (Step width)

#### Replication

##### R1

We found very strong evidence (BF_+0_ = 37.47) that a perturbing force field results in an increased degree of foot placement control as an after-effect (R1, Figure SB1).

**Figure SB 1.**
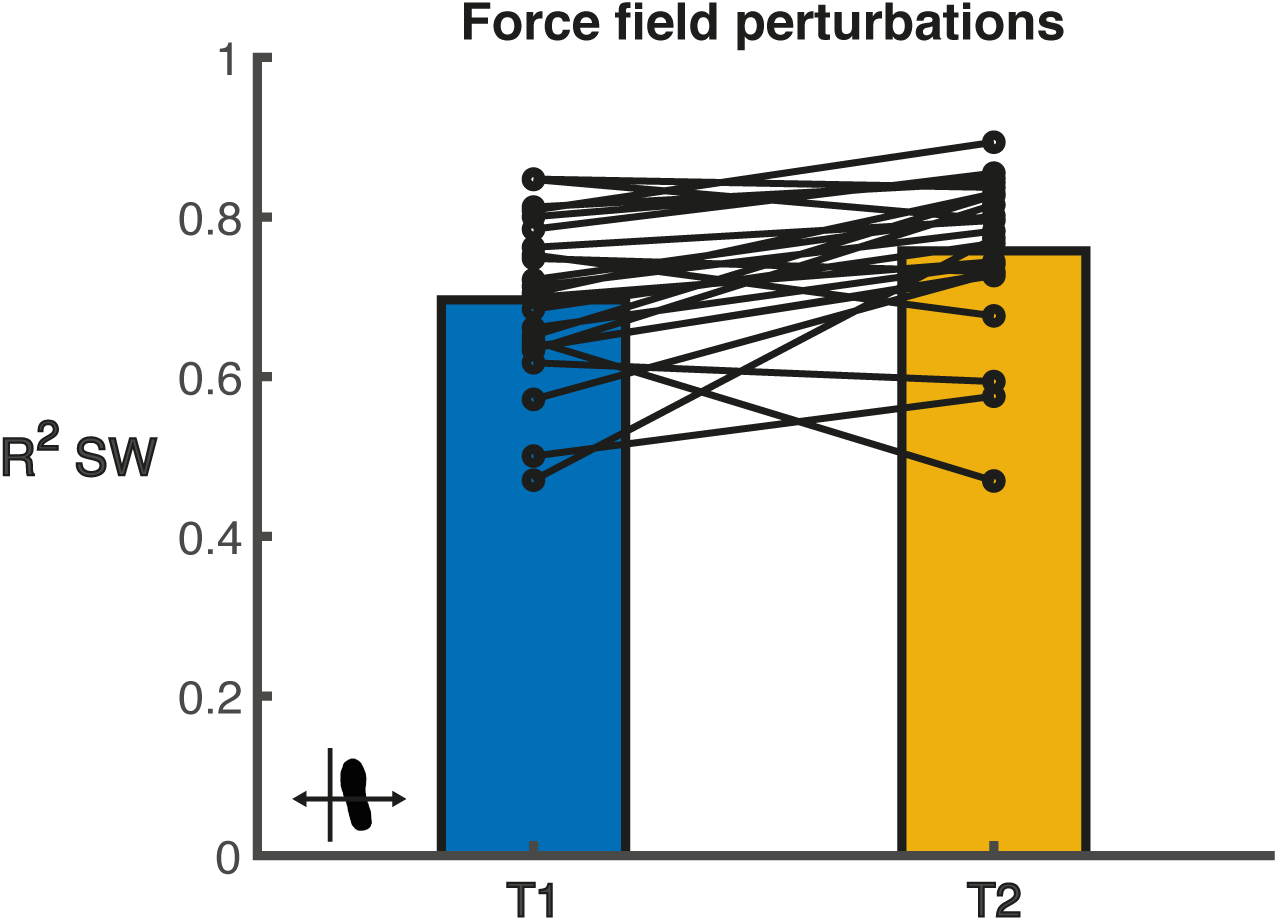
The degree of foot placement control (R^2^) before (T1) and after (T2) force field perturbations. The black circles indicate individual data points. R^2^ was determined from the foot placement model with SW as the dependent variable and CoM kinematic state predictors at terminal swing.

Foot placement error (extreme evidence, BF_10_ = 143), variability in mediolateral CoM position (moderate evidence, BF_10_ = 9.83) and the regression coefficient of ML_CoM_ (very strong evidence, BF_10_ = 90.58) all increased as an after-effect.

**Figure SB 2.**
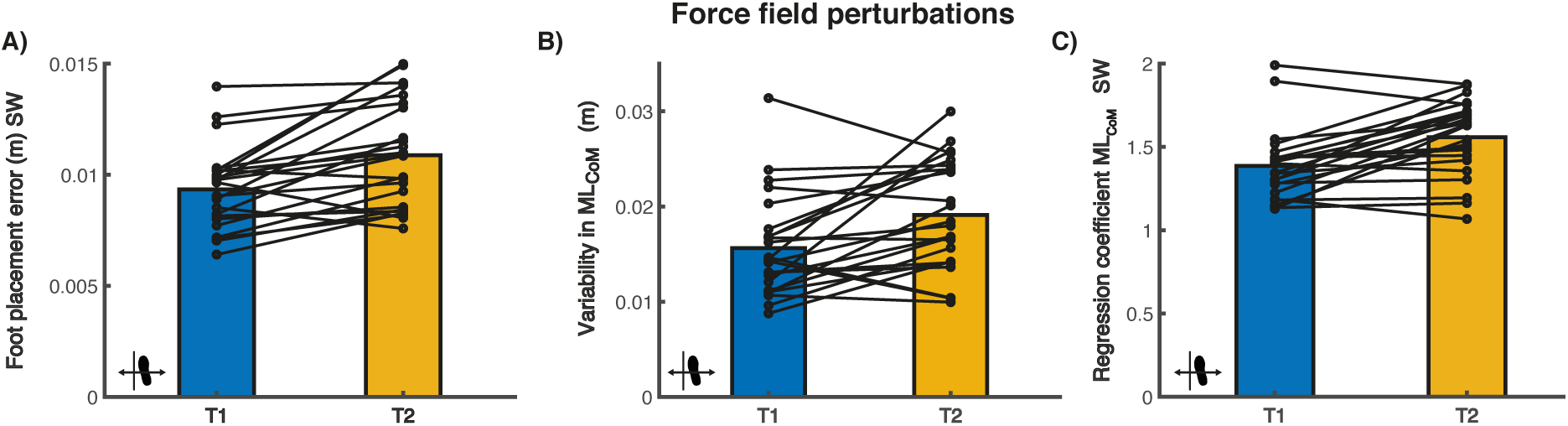
Foot placement error (Panel A), mediolateral CoM position variability (panel B) and β_pos_ (panel C) before (T1) and after (T2) force field perturbations. The black circles indicate individual data points. These secondary outcomes provide more insight into the underlying foot placement control changes reflected in the R^2^ of figure SB 1.

##### R2

We found extreme evidence, (BF_+0_ = 466.67) that augmented proprioception enhances the degree of foot placement control during the muscle vibration (R2, Figure SB3).

**Figure SB 3.**
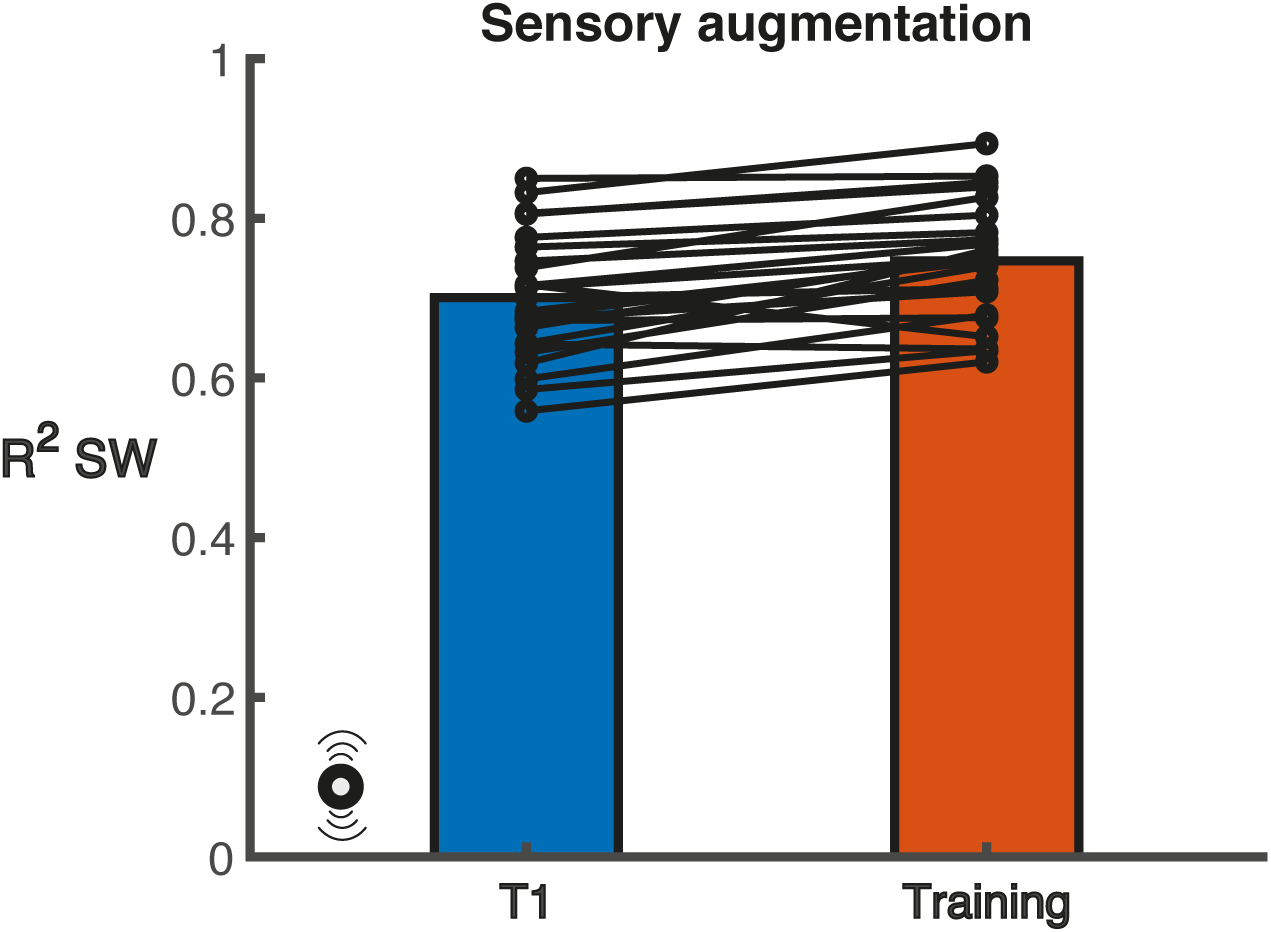
The degree of foot placement control (R^2^) before (T1) and during (Training) sensory augmentation. The black circles indicate individual data points. R^2^ was determined from the foot placement model with SW as the dependent variable and CoM kinematic state predictors at terminal swing.

**Figure SB4.**
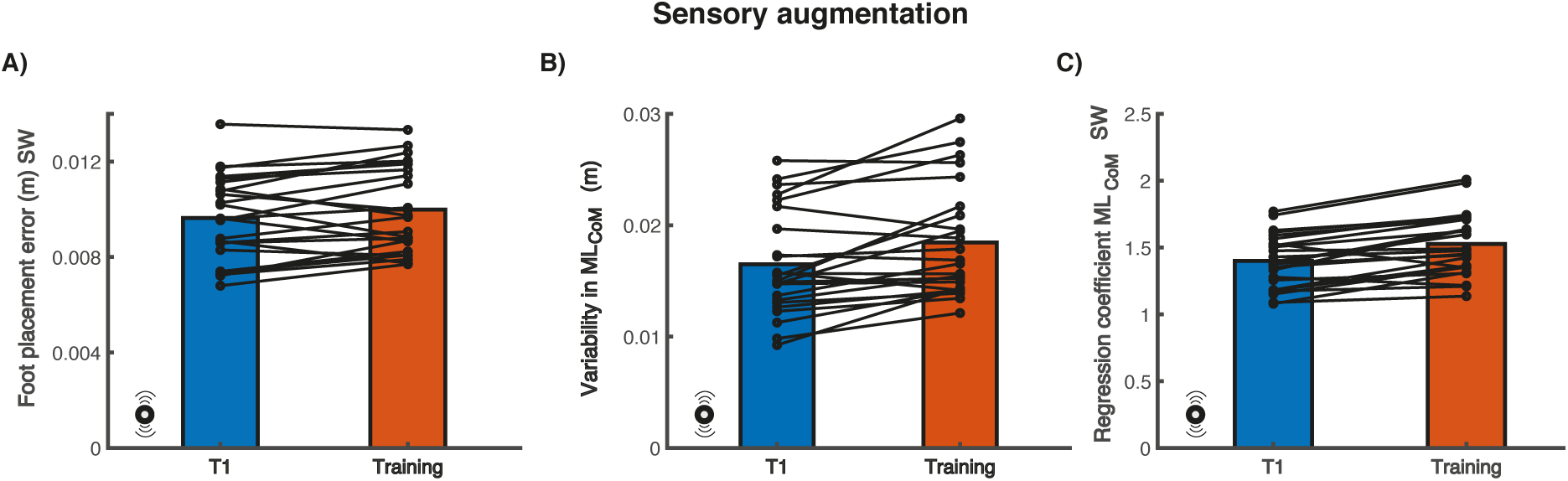
Foot placement error (Panel A), mediolateral CoM position variability (panel B) and β_pos_ (panel C) before (T1) and during (Training) sensory augmentation. The black circles indicate individual data points. These secondary outcomes provide more insight into the underlying foot placement control changes reflected in the R^2^ of figure SB3.

### Q1: Sensory augmentation after-effect?

We found extreme evidence (BF_10_ = 233.36) that sensory augmentation caused an after-effect (Q1). In Figure SB 5 it can be seen that the degree of foot placement control increased in T2 as compared to T1.

**Figure SB 5.**
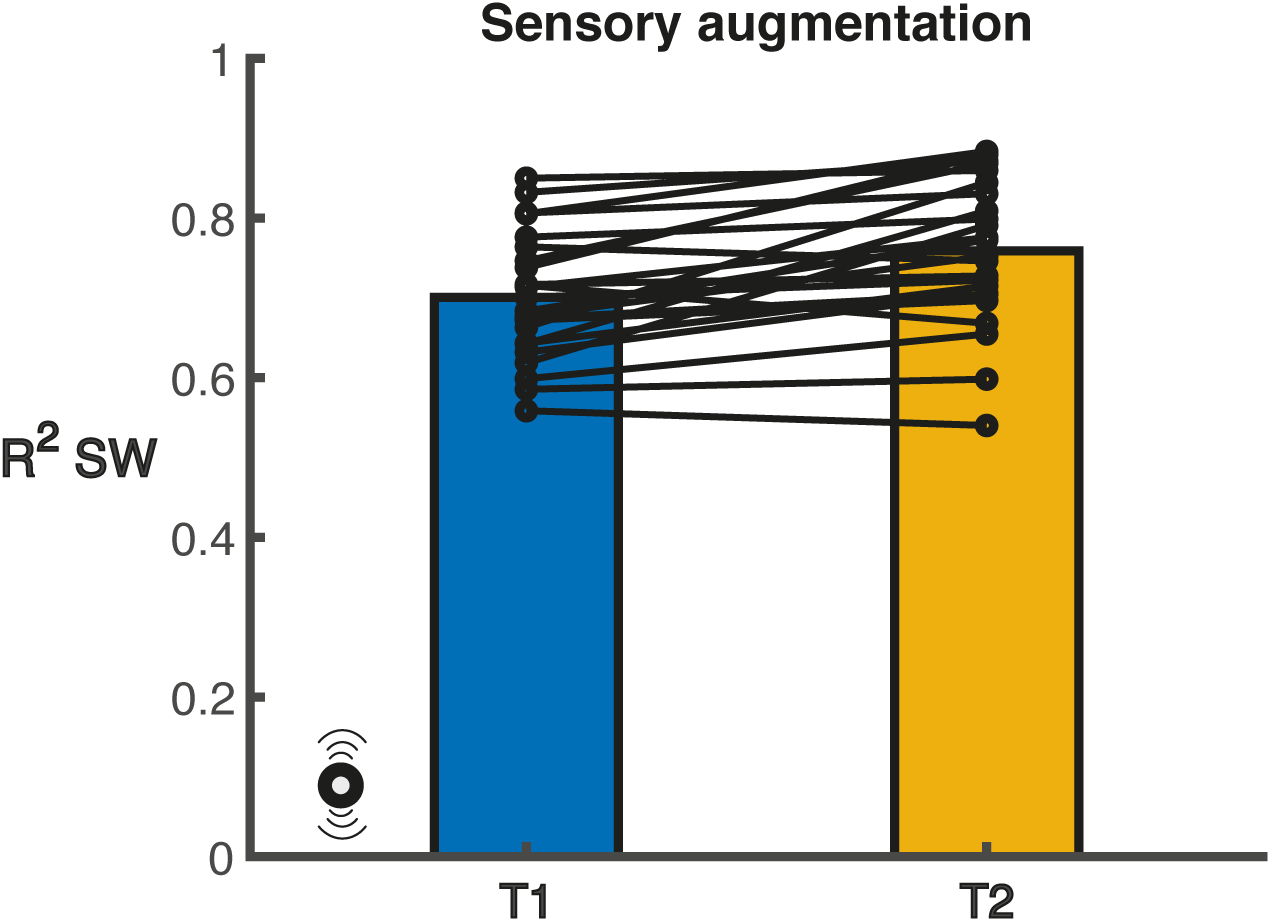
The degree of foot placement control (R^2^) before (T1) and after (T2) sensory augmentation. The black circles indicate individual data points. R^2^ was determined from the foot placement model with SW as the dependent variable and CoM kinematic state predictors at terminal swing.

**Figure SB 6.**
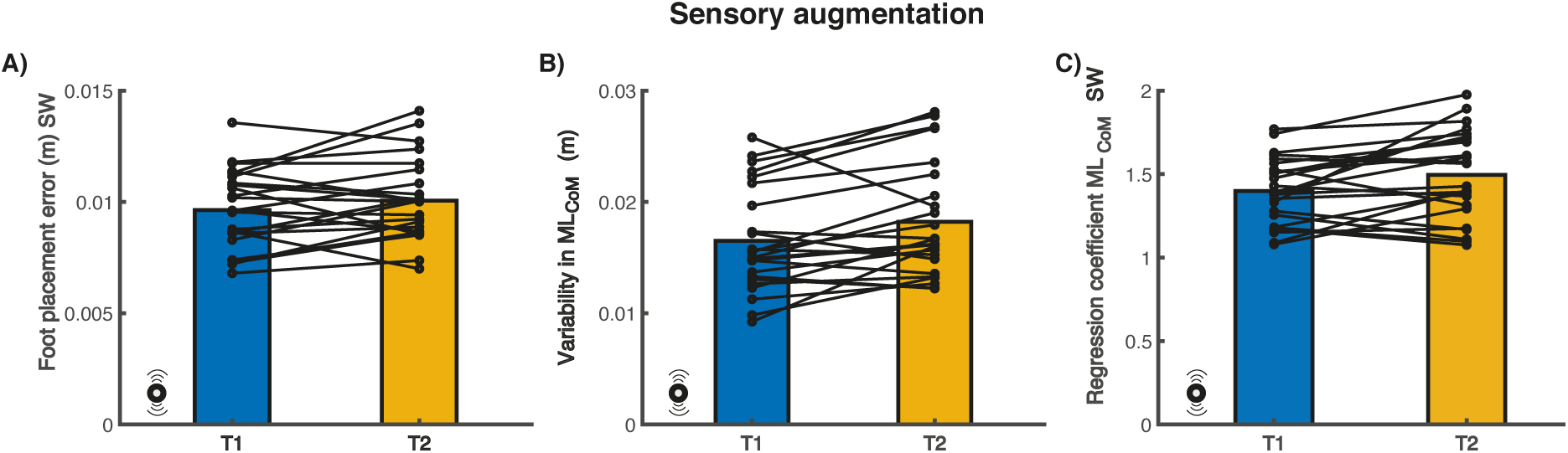
Foot placement error (Panel A), mediolateral CoM position variability (panel B) and β_pos_ (panel C) before (T1) and after (T2) sensory augmentation. The black circles indicate individual data points. These secondary outcomes provide more insight into the underlying foot placement control changes reflected in the R^2^ of figure SB5.

### Force-field perturbations and sensory augmentation

**Figure SB 7.**
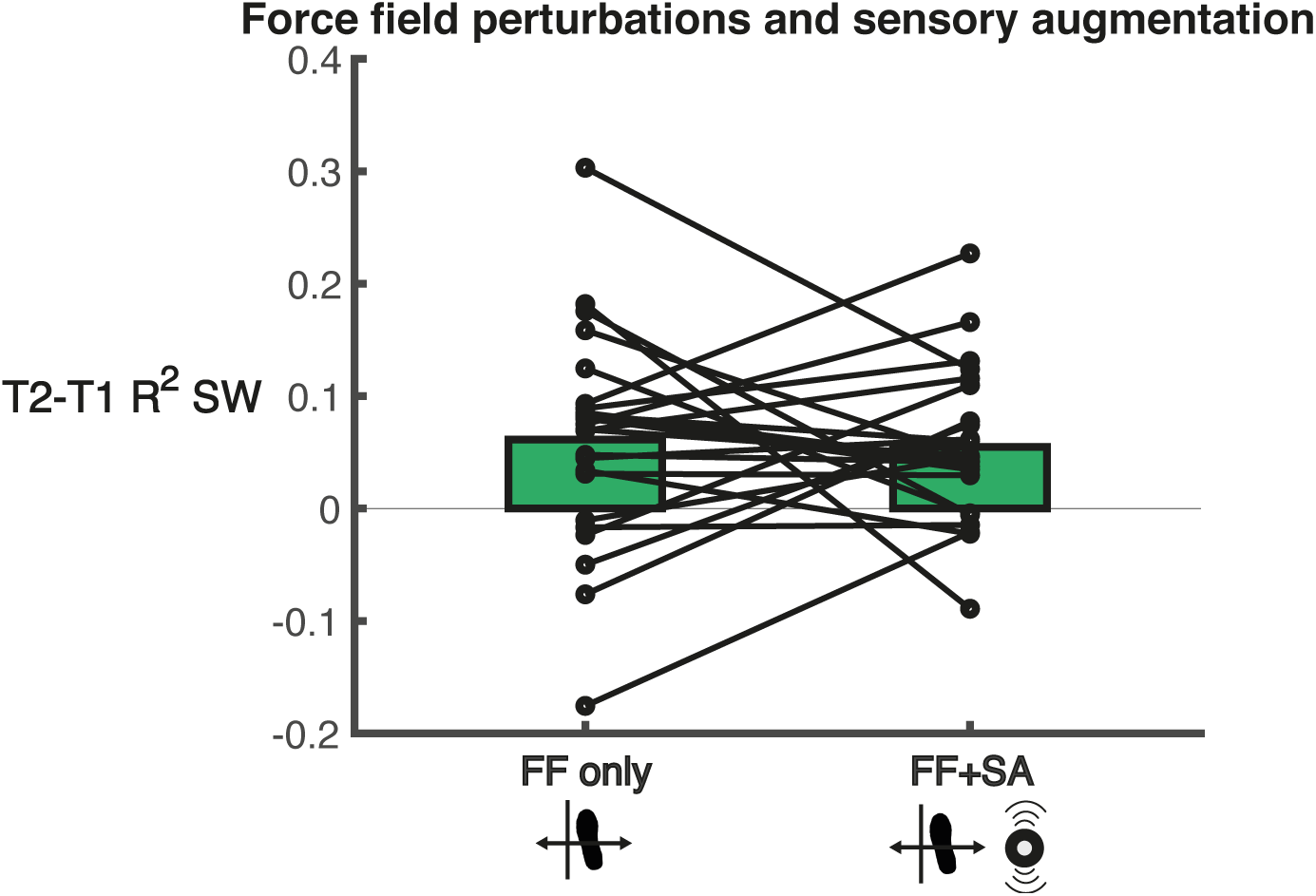
The change in the degree of foot placement control (R^2^) between T2 and T1 after force-field perturbations only and after a combination of force-field perturbations and sensory augmentation. The black circles indicate individual data points. R^2^ was determined from the foot placement model with SW as the dependent variable and CoM kinematic state predictors at terminal swing.

**Figure SB 8.**
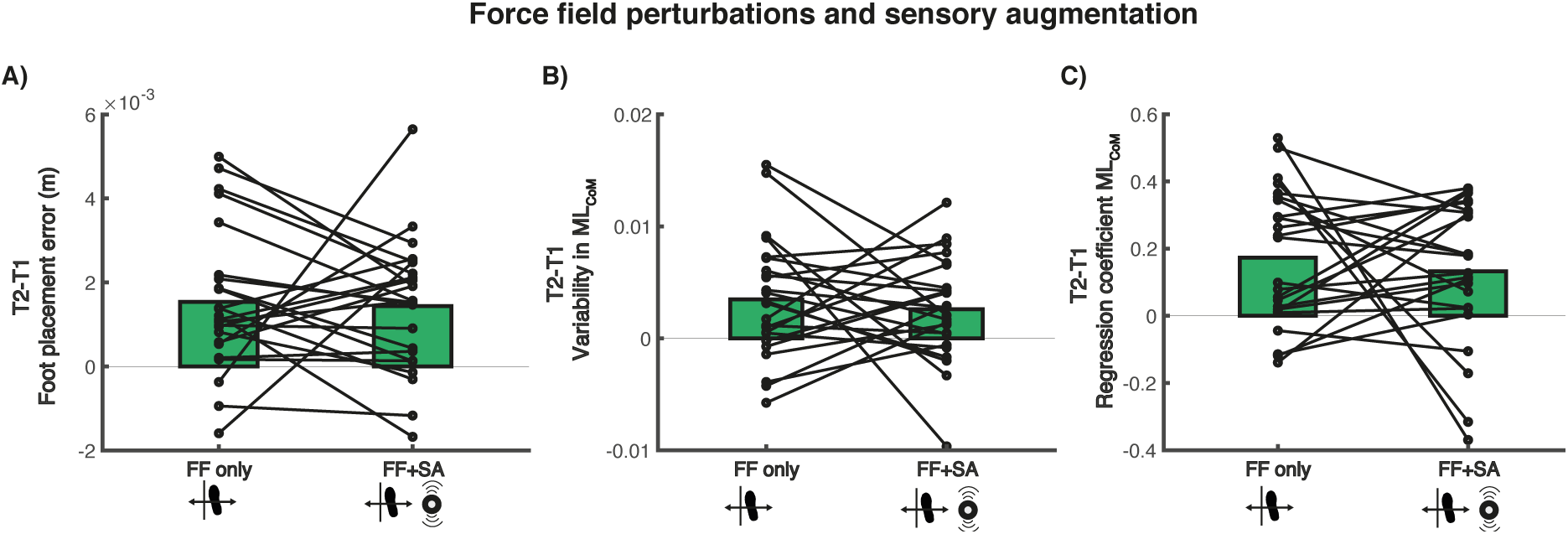
The change in foot placement error (Panel A), mediolateral CoM position variability (panel B) and β_pos_ (panel C), between T2 and T1 after force-field perturbations only and after a combination of force-field perturbations and sensory augmentation. The black circles indicate individual data points. These secondary outcomes provide more insight into the underlying foot placement control changes reflected in the delta R^2^ of figure 10.

## References

1. Bruijn, S.M. and J.H. van Dieën, Control of human gait stability through foot placement. Journal of The Royal Society Interface, 2018. 15(143): p. 20170816.

2. Hof, A.L., The equations of motion for a standing human reveal three mechanisms for balance. Journal of biomechanics, 2007. 40(2): p. 451–457.

3. Rankin, B.L., S.K. Buffo, and J.C. Dean, A neuromechanical strategy for mediolateral foot placement in walking humans. Journal of neurophysiology, 2014. 112(2): p. 374–383.

4. van Leeuwen, A.M., et al., Active foot placement control ensures stable gait: Effect of constraints on foot placement and ankle moments. Plos one, 2020. 15(12): p. e0242215.

5. Wang, Y. and M. Srinivasan, Stepping in the direction of the fall: the next foot placement can be predicted from current upper body state in steady-state walking. Biology letters, 2014. 10(9).

6. Hurt, C.P., et al., Variation in trunk kinematics influences variation in step width during treadmill walking by older and younger adults. Gait & posture, 2010. 31(4): p. 461–464.

7. Arvin, M., et al., Where to step? Contributions of stance leg muscle spindle afference to planning of mediolateral foot placement for balance control in young and older adults. Frontiers in physiology, 2018. 9: p. 1134.

8. Stimpson, K.H., et al., Post-stroke deficits in the step-by-step control of paretic step width. Gait & posture, 2019. 70: p. 136–140.

9. Dean, J.C. and S.A. Kautz, Foot placement control and gait instability among people with stroke. Journal of rehabilitation research and development, 2015. 52(5): p. 577.

10. Mahaki, M., S.M. Bruijn, and J.H. Van Dieën, The effect of external lateral stabilization on the use of foot placement to control mediolateral stability in walking and running. PeerJ, 2019. 7: p. e7939.

11. van Leeuwen, A., J. van Dieën, and S. Bruijn, The effect of external lateral stabilization on ankle moment control during steady-state walking. bioRxiv, 2022.

12. Reimold, N.K., et al., Effects of targeted assistance and perturbations on the relationship between pelvis motion and step width in people with chronic stroke. IEEE Transactions on Neural Systems and Rehabilitation Engineering, 2020. 29: p. 134–143.

13. Reimold, N.K., et al., Altered active control of step width in response to mediolateral leg perturbations while walking. Scientific reports, 2020. 10(1): p. 1–15.

14. Heitkamp, L.N., K.H. Stimpson, and J.C. Dean, Application of a novel force-field to manipulate the relationship between pelvis motion and step width in human walking. IEEE Transactions on Neural Systems and Rehabilitation Engineering, 2019. 27(10): p. 2051–2058.

15. van Leeuwen, A., et al., Ankle muscles drive mediolateral center of pressure control to ensure stable steady state gait. Scientific reports, 2021. 11(1): p. 1–14.

16. Bastian, A.J., Understanding sensorimotor adaptation and learning for rehabilitation. Current opinion in neurology, 2008. 21(6): p. 628.

17. Knapp, H.A., B.A. Sobolewski, and J.C. Dean, Augmented hip proprioception influences mediolateral foot placement during walking. IEEE Transactions on Neural Systems and Rehabilitation Engineering, 2021. 29: p. 2017–2026.

18. Nyberg, E.T., et al., A novel elastic force-field to influence mediolateral foot placement during walking. IEEE Transactions on Neural Systems and Rehabilitation Engineering, 2016. 25(9): p. 1481–1488.

19. Zeni Jr, J., J. Richards, and J. Higginson, Two simple methods for determining gait events during treadmill and overground walking using kinematic data. Gait & posture, 2008. 27(4): p. 710–714.

20. Mahaki, M., et al., *Foot placement control can be trained:* Older adults learn to walk more stable, when ankle moments are constrained. bioRxiv, 2023: p. 2023.03.10.532038.

21. Hoogstad, L.A., et al., Can foot placement during gait be trained? Adaptations in stability control when ankle moments are constrained. Journal of Biomechanics, 2022: p. 110990.

22. Kelter, R., Bayesian alternatives to null hypothesis significance testing in biomedical research: a non-technical introduction to Bayesian inference with JASP. BMC Medical Research Methodology, 2020. 20(1): p. 1–12.

23. Sansare, A., et al., Subthreshold electrical noise alters walking balance control in individuals with cerebral palsy. Gait & Posture, 2023. 106: p. 47–52.

24. Meyer, C., et al., Familiarization with treadmill walking: How much is enough? Scientific reports, 2019. 9(1): p. 1–10.

25. Howard, K.E., et al., Relationships between mediolateral step modulation and clinical balance measures in people with chronic stroke. bioRxiv, 2022: p. 2022.04.26.489530.

